# sBOSC: A method for source-level identification of neural oscillations in electromagnetic brain signals

**DOI:** 10.1101/2025.07.20.665618

**Authors:** Enrique Stern, Guiomar Niso, Almudena Capilla

## Abstract

Neural oscillations are recognized as a fundamental component of brain electromagnetic activity. They are implicated in a wide range of cognitive processes and proposed as a core mechanism for brain communication. Nonetheless, detecting genuine neural oscillations remains a methodological challenge, particularly due to the difficulty of distinguishing them from aperiodic background activity. To identify episodes of oscillatory activity directly at their sources, we developed sBOSC, which extends the BOSC (Better OSCillation detection) family of algorithms. Consistent with existing approaches, sBOSC detects oscillatory episodes that exceed both a defined power threshold and a minimum duration criterion. In sBOSC, however, the detection of oscillatory episodes also relies on identifying peaks (i.e., local maxima) in the power spectra as well as throughout the brain volume (spatial peaks). Using a series of simulated signals, we tested the ability of sBOSC to detect and localize oscillations across multiple scenarios. Our results show that most oscillatory episodes were accurately detected at their sources, achieving above 95% accuracy under optimal conditions (i.e., high signal-to-noise ratio, lower frequencies, and numerous successive cycles). In addition, we validated sBOSC’s performance using real magnetoencephalography (MEG) data from both resting-state and motor task recordings. From the detected oscillatory episodes, we extracted a topographic distribution of natural frequencies that is consistent with previous work, as well as the expected alpha- and beta-band modulations over sensorimotor regions during motor preparation. In conclusion, sBOSC offers a novel approach for identifying oscillatory activity in electrophysiological signals. It extends previous algorithms by operating in source space and verifying the presence of genuine spectral peaks, thereby enabling new possibilities for exploring brain dynamics.

## 1. Introduction

Neural oscillations have become a major area of interest in cognitive neuroscience, as different frequencies have been linked to a wide variety of cognitive processes and functions (Cannon et al., 2014; Helfrich et al., 2018; Herweg et al., 2020; Lega et al., 2012; Noah et al., 2020; Shin et al., 2017). Furthermore, several theoretical frameworks have posited brain oscillations as a fundamental mechanism for neural communication and temporal organization of brain function (Bonnefond et al., 2017; Buzsáki & Watson, 2012; Fries, 2015; Izhikevich et al., 2003; Palva & Palva, 2018; Vinck et al., 2023).

Nonetheless, studying neural oscillations constitutes a challenging task, since they are obscured by the so-called aperiodic component, a broadband signal that follows a power-law function, where power decreases exponentially with increasing frequency (1/f^β^). This component is also referred to as arrhythmic, scale-free, pink noise, or background activity (He, 2014), indicating the lack of any characteristic frequency in the power spectrum. The aperiodic component has been suggested to reflect key physiological processes such as the balance between excitation and inhibition in neural circuits (Gao et al., 2017) and the population fire rate (He, 2014) and has been associated with task performance and various clinical conditions (Donoghue, 2024).

While most studies have traditionally relied on baseline subtraction to attenuate the aperiodic component (Gross, 2014; Gyurkovics et al., 2022), the idea of parameterizing the power spectrum to isolate oscillatory activity is rising in popularity (Donoghue et al., 2020). Thus, to confirm the presence of a neural oscillation, a narrow-band frequency peak must be detected over the aperiodic component (Donoghue et al., 2022; Gross, 2014; Lopes da Silva, 2013). Rhythmicity has also been proposed as a key feature for detecting brain oscillatory activity (Fransen et al., 2015), although its temporal manifestation is currently debated, since oscillations do not always manifest as sustained patterns, but often occur in the form of transient bursts (Jones, 2016; Van Ede et al., 2018; Vidaurre et al., 2016).

Driven by a growing recognition of their theoretical and methodological relevance, recent years have witnessed a surge in methods for detecting transient episodes of oscillatory brain activity in electrophysiological data. A family of algorithms designed for this purpose is BOSC (Better OSCillation detection). A common practice across the different BOSC variants is to establish both a duration and a power threshold. The duration threshold is typically set at three consecutive cycles to minimize spurious results and guarantee rhythmicity. The definition of the power threshold has evolved with successive versions of the method. For example, in its original formulation (BOSC; Caplan et al., 2001; Whitten et al., 2011), a linear model of the aperiodic component was used to estimate the power threshold corresponding to the 95^th^ percentile of a χ^2^ distribution. The method was later extended (eBOSC; Kosciessa et al., 2020) and adapted to fit the model using robust regression, thus improving specificity of oscillatory activity detection. More recently, FOOOF (Fitting Oscillations and One-Over-F; Donoghue et al., 2020), a popular spectral parameterization algorithm, has been incorporated within the BOSC framework to improve the estimation of the aperiodic component (fBOSC; Seymour et al., 2022). A related method outside the BOSC framework is PAPTO (Periodic/Aperiodic Parameterization of Transient Oscillations; Brady & Bardouille, 2022), which also uses FOOOF to model the aperiodic background and sets the power threshold as a multiple of the modelled aperiodic component.

While oscillation detection methods have advanced considerably, two main challenges remain. First, as noted above, oscillations must manifest as narrow-band peaks in the power spectrum (Donoghue et al., 2022; Gross, 2014; Lopes da Silva, 2013). Within the family of BOSC algorithms, all frequency bins exceeding the power threshold are considered potential oscillations. However, it is only the frequency peak in the power spectrum that defines the underlying oscillation. This is of particular importance when multiple frequencies coexist (e.g., 10 and 20 Hz), as only their respective peaks indicate genuine oscillations, whereas intermediate frequencies (e.g., 15 Hz), even if they exceed the power threshold, do not reflect true oscillatory activity. In practice, these intermediate frequencies rarely exceed the duration threshold and are therefore usually not selected as episodes. Additionally, eBOSC incorporates controls for wavelet-induced frequency and temporal leakage, which further mitigates this problem. Nevertheless, plateaus across these frequencies can still occasionally be erroneously categorized as independent oscillations, a problem that can be addressed by detecting local maxima in the power spectrum.

The second challenge is the identification of oscillatory episodes across different brain regions. Existing methods are typically limited to single-channel electrophysiological signals, yet a comprehensive characterization of brain oscillations requires incorporating multi-channel information, as provided by magneto-/electroencephalography (M/EEG) recordings. Furthermore, the identification of oscillations at the sensor level does not necessarily imply the presence of an underlying neural generator, since rhythmic fluctuations in the M/EEG signal can arise from multiple factors (Cohen, 2017; Jackson & Bolger, 2014; Van Bree et al., 2024). In contrast, detecting oscillations directly at the source level would enable the identification of the underlying oscillatory generators, thereby improving both the spatial precision of detection and the interpretability of the results.

To address these challenges, we have developed an algorithm within the BOSC framework to detect episodes of brain oscillatory activity at the source level in non-invasive, whole-brain electrophysiological data (M/EEG). The method, called sBOSC for source-BOSC, incorporates several improvements. First, to ensure the detection of genuine oscillatory activity, only peaks in the power spectrum are selected. Second, we only take into consideration oscillatory activity in voxels presenting a spatial peak in oscillatory power, thus mitigating undesired source leakage effects. In this study, we validated sBOSC by applying it to a series of simulated signals with varying parameters, such as oscillatory frequency, number of cycles, generator depth, and signal-to-noise ratio (SNR). Additionally, sBOSC was applied to empirical MEG data to evaluate its effectiveness in real-world scenarios. We first applied sBOSC to resting-state MEG data to assess whether the natural frequencies (i.e., the most characteristic oscillatory frequencies at each voxel) estimated using this method were consistent with those reported in previous studies (Arana et al., 2025; Capilla et al., 2022). Similarly, we evaluated the performance of sBOSC on task-related MEG data from a hand movement preparation task, which is well known to modulate alpha- and beta-band oscillations over sensorimotor regions (Pfurtscheller & Lopes da Silva, 1999; Salmelin & Hari, 1994).

## 2. Methods

Here, we have developed sBOSC, a method for detecting oscillatory episodes in source space from electrophysiological brain signals. As in previous BOSC variants, oscillatory episodes were defined as segments of the signal that exceed both a power threshold and a duration threshold. The power threshold was set at the 95th percentile of the power of the aperiodic component. Furthermore, only peaks identified in both the power spectrum and the brain volume were retained to ensure the selection of genuine oscillatory activity. The duration threshold required a minimum of three complete cycles to confirm rhythmicity.

To validate sBOSC, we applied it to simulated MEG data, which provide a known ground truth and allow for a precise and quantitative evaluation of the method’s performance under realistic MEG conditions. To ensure its applicability and effectiveness in real-world scenarios, we then applied sBOSC to empirical MEG data, including recordings from both resting-state and a motor task. The following sections describe the generation of the simulated data and the acquisition procedures for real MEG recordings. We then provide a detailed description of the sBOSC algorithm, followed by an explanation of how datasets were analyzed to evaluate the performance and validity of the method. All code was written in MATLAB (2024b; The MathWorks Inc, 2024), with additional functions from FieldTrip (version 20230118; Oostenveld et al., 2011), fBOSC (Seymour et al., 2022), and FOOOF (Donoghue et al., 2020). The sBOSC toolbox and the full code needed to reproduce the analyses and figures in this work are publicly available at https://github.com/necog-UAM/.

### 2.1. Generation of simulated data

To evaluate the performance of sBOSC, we simulated brain oscillatory activity embedded in realistic background noise (Figure 1). Data from a randomly selected participant (see section 2.4) were used as a template to generate both a simulated brain volume and the MEG sensor coordinates. A pure sinusoidal oscillation was first generated at a specific voxel and time window and then projected onto the sensor space. To mimic the coexistence of oscillatory and aperiodic components in real electrophysiological recordings, pink noise—a hallmark of aperiodic activity—was generated at five semi-random brain locations using the *cnoise* function implemented in fBOSC (Seymour et al., 2022). Candidate voxels for aperiodic sources were constrained to lie at least 1.5 cm away from oscillatory sources, thus minimizing contamination and ensuring the spatial specificity of oscillation localization. Aperiodic activity was subsequently projected onto the sensors. Additionally, white noise with an amplitude equivalent to the median of the aperiodic activity was added to account for stochastic noise inherent to MEG recordings. Finally, the oscillatory, aperiodic and white noise components were linearly combined to obtain the simulated signal.

**Figure 1.**
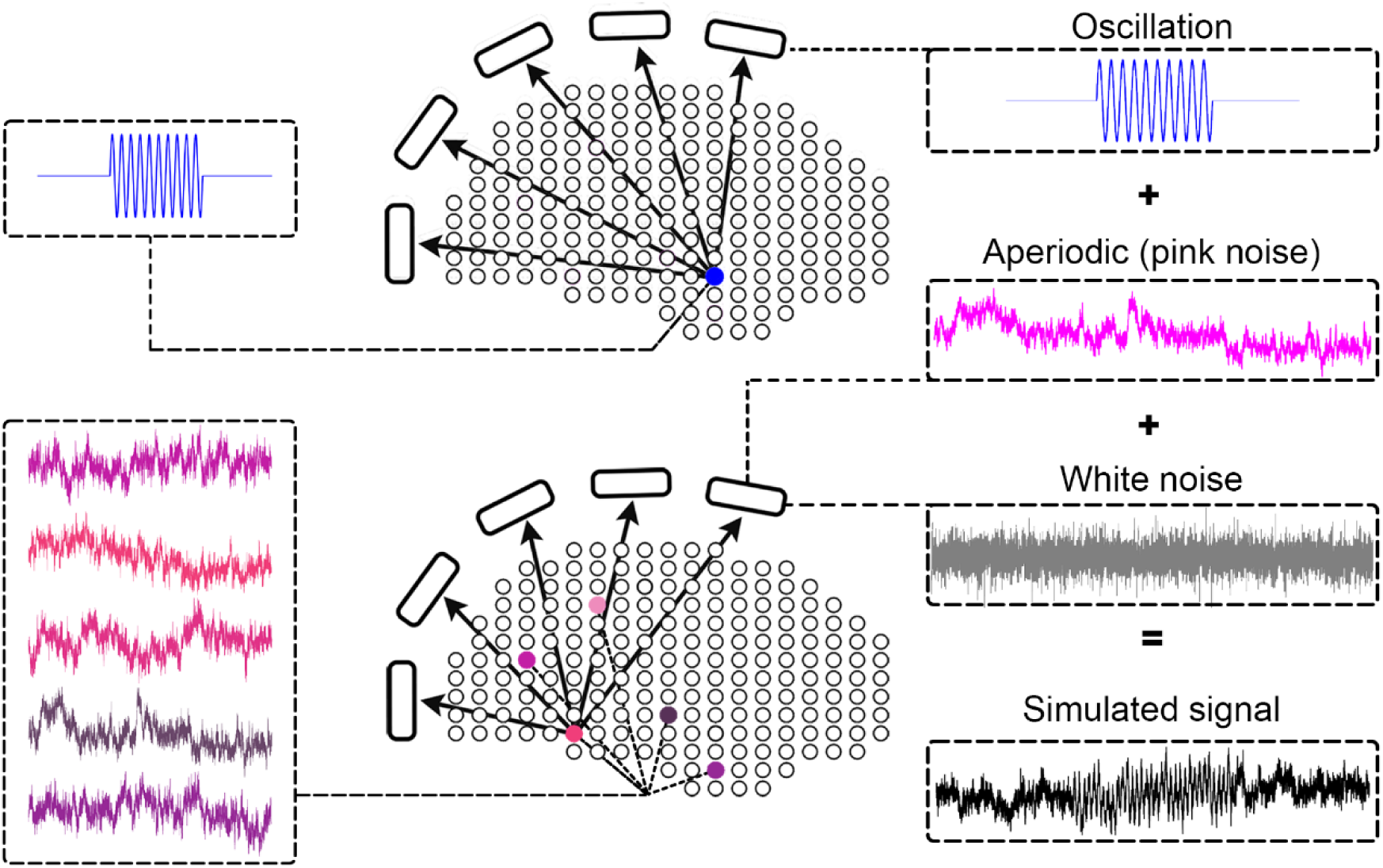
Simulation of brain signals in MEG sensors. A sinusoidal oscillation was generated at a specific voxel in the brain and projected onto the sensors (blue). Pink noise was generated at five semi-randomly selected voxels and subsequently projected onto the sensors (pink). Oscillatory activity, pink noise, and white noise (gray) were linearly combined to create the simulated brain signal in sensor space (black).

We simulated multiple datasets by systematically varying different parameters that could potentially influence the detection performance of sBOSC: oscillatory frequency (5, 10, and 20 Hz), number of cycles (3, 10, and 20 consecutive cycles), generator depth (65, 75, and 85 mm from the MEG sensors), and SNR (0.5, 1.0, and 1.5). SNR was computed as the ratio between the root mean square (RMS) of the sinusoidal oscillation and the RMS of the pink noise in the sensor exhibiting the strongest projection of oscillatory activity. In addition, to ensure that the spatial configuration of the aperiodic sources did not affect sBOSC performance, the simulation procedure was repeated five times for each parameter combination, using a new set of semi-randomly selected pink noise voxels in each repetition. Therefore, for each of the four parameters (e.g., oscillatory frequency), we generated three different signals (e.g., 5, 10, and 20 Hz), and repeated the process five times, resulting in a total of 405 simulated signals (3^4^ × 5).

### 2.2. Acquisition and pre-processing of resting-state MEG data

Anonymized resting-state MEG recordings and T1-weighted Magnetic Resonances Images (MRIs) were downloaded from the open-access OMEGA database (Niso et al., 2016). Data acquisition took place at the Montreal Neurological Institute (MNI, McGill University), with approval from the institutional ethics committees and in accordance with the declaration of Helsinki. MEG recordings were acquired using a whole-head CTF MEG system equipped with 275 axial gradiometers and 26 reference sensors, within a magnetically shielded room. The MEG signal was digitized at a sampling rate of 2400 Hz and recorded with a 600 Hz low-pass anti-aliasing filter. Individual head shapes and three anatomical landmarks (nasion, left- and right- pre-auricular points) were also digitized to facilitate MEG-MRI co-registration.

We analyzed data from 128 healthy participants (68 males, 118 right-handed, mean age 30.5 ± 12.4 [M ± SD], range 19-73 years) who were sitting upright and instructed to maintain a relaxed wakeful state with their eyes open, fixating on a cross for five minutes. In this study, we used data that had been preprocessed in a previous work (Capilla et al., 2022). In brief, MEG signals were denoised and filtered with a 0.05 Hz high-pass filter. Power line noise and its harmonics were reduced using spectrum interpolation. Finally, artifactual components related to cardiac, ocular, and muscular activity were removed using Independent Component Analysis (ICA), and contaminated data segments were excluded from further analysis.

### 2.3. Acquisition and preprocessing of motor MEG data

Similarly, anonymized MEG recordings and T1-weighted MRIs during the execution of a motor task were downloaded from the open-access Human Connectome Project database (HCP; WU-Minn Consortium Human Connectome Project, 2017; Larson-Prior et al., 2013; Van Essen et al., 2013). Data acquisition took place at the Saint Louis University medical campus, with approval from the institutional ethics committees and in consonance with the Declaration of Helsinki. MEG recordings were acquired in a magnetically shielded room using a whole-head MAGNES 3600 system (4D Neuroimaging, San Diego, CA) equipped with 248 magnetometers and 23 reference sensors. Recordings were obtained at ∼2034 Hz with a 400 Hz bandwidth.

We analyzed preproccesed, artifact-free data directly obtained from the HCP database from 61 participants (28 males, mean age 27.1 ± 3.8 [M ± SD], range 22-35 years). The preprocessing procedure included identification and removal of bad segments and bad channels, as well as artifact correction via ICA.

Briefly, the motor task consisted of a visually cued paradigm adapted from previous protocols (Buckner et al., 2011; Yeo et al., 2011). Participants performed repetitive movements—either tapping the thumb and index finger or flexing the toes—guided by 3-second visual cues indicating left or right hand or foot. For the purpose of validating sBOSC, we analyzed only left- and right-hand trials. To specifically examine the phase of movement preparation, the signal was segmented into trials starting 1.2 seconds before movement onset and ending at movement onset, as determined by the electromyography (EMG) signal.

### 2.4. Detection of oscillatory episodes using sBOSC

The sBOSC algorithm comprises the following eight steps: (1) reconstruction of source-level activity, (2) estimation of the aperiodic component, (3) correction of the center of the head bias, (4) time-frequency decomposition, (5) identification of the power threshold, (6) detection of peaks in the power spectra, (7) detection of spatial peaks in the brain volume, and (8) selection of oscillatory episodes exceeding the duration threshold. These analysis steps are illustrated in Figure 2 and explained in more detail in the next paragraphs.

**Figure 2.**
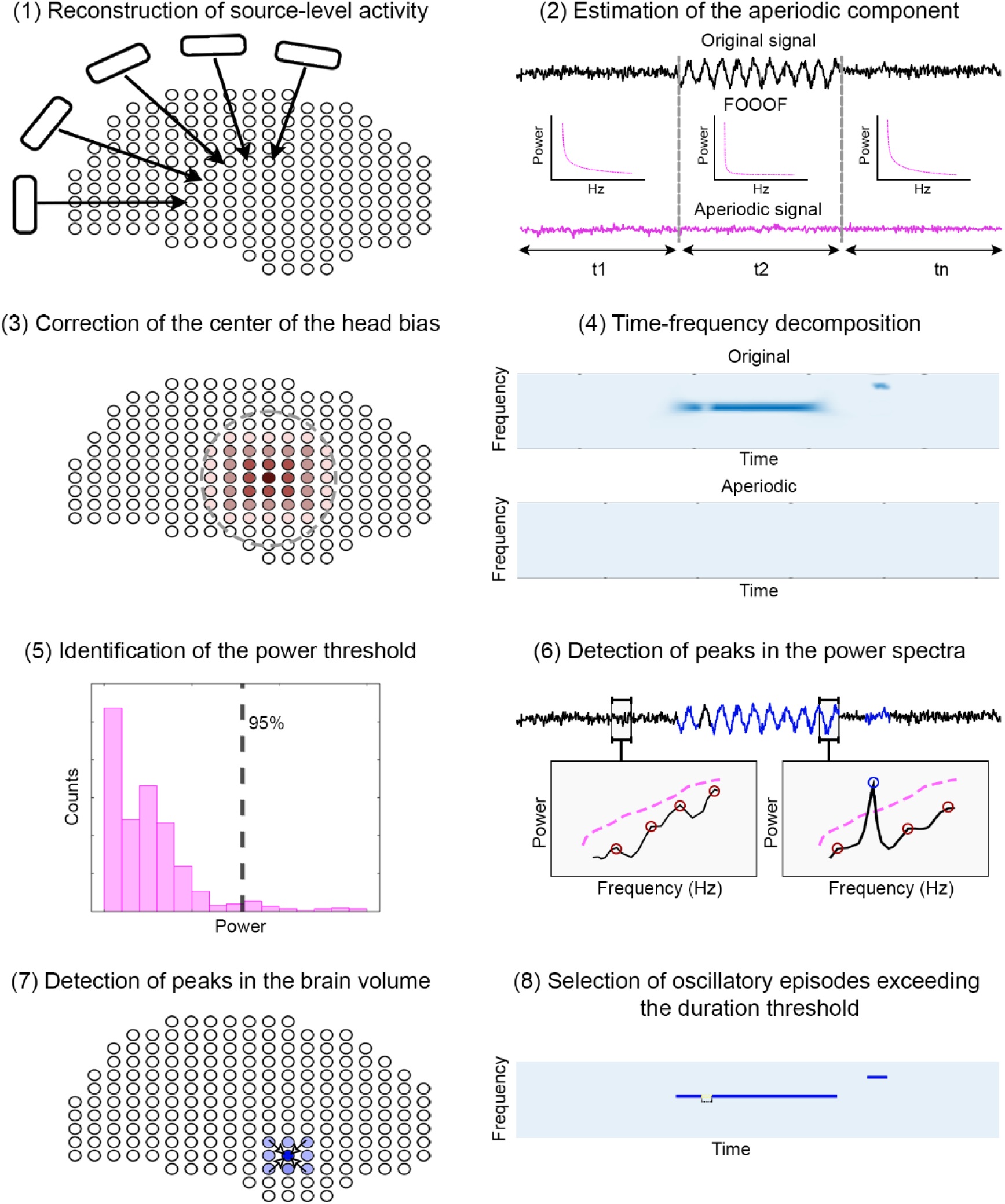
Detection of oscillatory episodes using sBOSC. **Step 1.** The pre-processed MEG signal is reconstructed in source space using beamforming. **Step 2.** The aperiodic component is extracted from the signal using FOOOF in 20-second time windows and transformed back to time domain. **Step 3.** The root mean square (RMS) of the aperiodic component is used to remove the center of the head bias introduced by beamformers. **Step 4.** Time-frequency decompositions of the source-reconstructed signal and the signal containing only the aperiodic component are computed using STFT. **Step 5.** The 95^th^ value of the aperiodic component distribution is used as a power threshold for each frequency. **Step 6.** Only time-frequency points that exceed the power threshold (dashed pink line) and display a peak in the power spectrum (red circles) are tagged (blue circle) as candidates for oscillations. Two illustrative spectra extracted are shown (boxes). **Step 7.** To mitigate spatial leakage effects, only local maxima in the brain volumes are considered oscillation generators. **Step 8.** Adjacent time-frequency points are connected to define oscillatory episodes, provided that at least three consecutive cycles are detected. A transiently discontinuous episode is highlighted and merged to form a continuous oscillatory episode. A false positive can be observed (right corner).

#### (1) Reconstruction of source-level activity

Since sBOSC has been designed to detect oscillatory activity directly in source space, the first step involves reconstructing the source-level time series from the MEG sensor data. For the resting-state MEG data, the co-registration of each participant’s T1-weighted MRI and the MEG head coordinate space was obtained from Capilla et al. (2022) (https://github.com/necog-UAM/OMEGA-NaturalFrequencies/tree/main/coregistration). Similarly, MRI-MEG co-registration for the motor MEG data was performed using the transformation matrices provided by the HCP Data Release (WU-Minn Consortium Human Connectome Project, 2017). For the simulated data, one randomly selected participant served as a template for defining the brain volume and MEG sensor locations used in every simulation. From this point onward, the analysis pipeline was identical for both the simulated and real data. A realistic single-shell volume conduction model (Nolte, 2003) was employed to compute the lead fields on a 1 cm resolution grid. Source-level time series were then reconstructed using linearly constrained minimum variance beamforming (LCMV; Van Veen et al., 1997). The covariance of the artifact-free data served to calculate the spatial filter weights, with the regularization parameter lambda adjusted to 10%. Lastly, beamforming weights were applied to reconstruct source-space time series from the sensor-level data.

#### (2) Estimation of the aperiodic component

Similarly to fBOSC and PAPTO, we estimated the aperiodic component from the source-reconstructed time series using the FOOOF algorithm (Donoghue et al., 2020), a popular method for parameterizing the aperiodic component of the signal. The estimation of the aperiodic activity served two purposes. First, it was employed to correct the center of the head bias, which is inherent in beamforming source reconstruction (step 3). Second, it provided the basis for determining the power threshold above which oscillations can be identified (step 5). Instead of computing a single aperiodic estimate for the entire signal, we applied FOOOF to shorter time windows, each lasting 20 seconds, to capture potential dynamic variations. For the analysis of the motor task, this window was adapted to match the duration of the trial (1.2 seconds). The estimate of the aperiodic component extracted from each time window was transformed back into the time domain. In this way, we obtained a signal containing only the aperiodic time series derived from FOOOF, subjected from that point onward to the same analysis procedures as the original signal.

#### (3) Correction of the center of the head bias

Beamformer source estimates are biased toward the center of the head, which exhibits stronger activity compared to cortical areas (Shapira Lots et al., 2016). This is typically corrected by normalizing the source-reconstructed activity with an estimate of noise. In Capilla et al. (2022), noise was estimated from the RMS of source-level data for each voxel. However, regions with high-amplitude oscillatory activity also tend to have high RMS values. For this reason, in this study, we estimated noise using the RMS of the signal containing only the aperiodic component, which, by definition, do not contain structured oscillatory activity. Thus, the center of the head bias was corrected in both the original and the aperiodic time series by normalizing them with the RMS of the aperiodic time series.

#### (4) Time-frequency decomposition

After downsampling the signal to 128 Hz to reduce computational load, a time-frequency decomposition was performed on both sets of data (original and signal containing only the aperiodic component) using the same parameters. Power values were extracted for 32 logarithmically spaced frequency bins, ranging from 1.8 to 40.5 Hz, using the Short-Time Fourier Transform (STFT) with a Hanning-tapered sliding window. The window length was adaptive, dynamically adjusted to cover 5 cycles of the frequency being analyzed. Previous BOSC variants have typically employed wavelet decomposition for time-frequency analysis. However, wavelets and STFT with a Hanning sliding window set to a fixed number of cycles per frequency produce largely equivalent results (Bruns, 2004).

#### (5) Identification of the power threshold

For each voxel and frequency, the power threshold was set to the 95^th^ percentile of the power derived from the signal containing only the aperiodic component across all time points. Thus, only time-frequency points exceeding this power threshold were considered potential candidates for oscillations in subsequent analysis.

#### (6) Detection of peaks in the power spectra

Since oscillations should exhibit a spectral peak (Donoghue et al., 2022; Gross, 2014; Lopes da Silva, 2013), local maxima were identified in the power spectra for every voxel and time point. Data points that fulfilled this criterion were tagged as candidates for oscillations.

#### (7) Detection of spatial peaks in the brain volume

Additionally, due to the ill-posed nature of the inverse source reconstruction problem, the so-called spatial or source leakage poses a significant challenge (O’Neill et al., 2015). In brief, while brain activity may originate from a single voxel, it can spread across a larger region of adjacent voxels, none of which are true generators. To reduce source leakage, we computed local maxima of spectral power spatially across the 3D brain volume and considered only the previously identified spectral peaks that also corresponded to a spatial peak within the brain.

#### (8) Selection of oscillatory episodes exceeding the duration threshold

Similarly to previous algorithms, we established a duration threshold to discard spurious and non-oscillatory transient brain events (Caplan et al., 2001; Kosciessa et al., 2020; Seymour et al., 2022). Thus, only episodes lasting at least three consecutive cycles were considered oscillatory episodes. To prevent false negatives due to brief power drops, sequential oscillatory bursts with the same frequency and appearing in the same voxel area (within a 1.5 cm radius) were merged if the temporal gap between them was less than half a cycle.

Importantly, although the analysis presented here used default parameters, the sBOSC toolbox allows users to modify several configuration options, enabling researchers to adapt the method to their specific needs. These options include applying the method to trial-based data, adjusting FOOOF input parameters, modifying the window length for time-frequency analysis, or defining the minimum number of consecutive cycles required for the duration threshold.

### 2.5. Analysis of simulated data

We applied sBOSC to the simulated data to evaluate its performance under controlled conditions. For each simulation, hit and false positive rates were quantified by averaging the outcomes across the five repetitions per parameter set. A hit, or correct detection, was defined as an episode detected at the same frequency and time points as simulated, and within a 1.5 cm radius of the true source location, thus accounting for slight inaccuracies in source reconstruction. False positives were defined as the total number of data points identified as episodes in the absence of simulated oscillations. To further examine the impact of the different parameters on sBOSC performance, we conducted the non-parametric version of the one-way analyses of variance (ANOVAs; Kruskal-Wallis) to compare the number of hits across all parameter settings (oscillatory frequency, number of cycles, generator depth, and SNR). Additionally, we verified that the semi-random location of the aperiodic generators did not affect the detection of oscillatory episodes.

To examine whether STFT window length might influence sBOSC’s performance, we additionally tested 3- and 7-cycle windows. The original simulation analyses were repeated 15 times with a constant SNR of 1 and the same oscillatory source. Frequency (5, 10, 20 Hz), duration (3, 5, 10 cycles), and window length (3, 5, 7 cycles) were systematically varied across repetitions.

### 2.6. Analysis of resting-state MEG data

To validate sBOSC under real-world conditions, we applied it to resting-state MEG data (Niso et al., 2016). This resulted in a matrix of detected oscillatory episodes, organized as time-points by frequencies, for each voxel within the cerebral cortex (1925 voxels) and for each participant (N = 128).

To compute the brain’s natural frequencies, representing its most characteristic oscillatory patterns, the total number of detected oscillatory cycles per participant, voxel, and frequency was summed across time. The resulting matrix was then z-score normalized across voxels to identify the frequency with the highest relative number of oscillatory cycles in each voxel. This normalization procedure aimed to minimize the dominance of the alpha rhythm (Capilla et al., 2022).

Natural frequencies at the group level were estimated following the bootstrapping procedure described by Capilla et al. (2022) to enable direct comparison with their results. In brief, participants were sampled with replacement 500 times. In each iteration, a histogram of the natural frequencies for each voxel was computed. Subsequently, we fitted a Gaussian function to all candidate peak frequencies and selected the peak with the highest goodness-of-fit. Each voxel’s natural frequency was then defined as the median value across bootstrap iterations. This method is preferred over the group mean, as some participants may have very extreme natural frequencies in the same voxel due to several reasons, including sharp transitions between nearby anatomical regions or limitations in the accuracy of source localization. Lastly, we validated our results by comparing the obtained map of natural frequencies with the one reported by Capilla et al. (2022). A Pearson correlation analysis was performed to assess the voxel-by-voxel similarity between the natural frequencies derived from both approaches. Additionally, for comparative purposes, we also computed the natural frequencies for 40 regions defined by the Automated Anatomical Labeling (AAL) atlas (Tzourio-Mazoyer et al., 2002).

### 2.7. Analysis of motor MEG data

Finally, we applied sBOSC to MEG data collected during a motor task to further extend its empirical validation to task-based experimental paradigms. We analyzed 1.2-second trials covering motor preparation until movement onset. For each individual trial, we extracted the average power and total duration of detected oscillatory episodes in the alpha and beta ranges. Both measures were respectively averaged across trials for each participant and subsequently spatially normalized by computing a z-score across voxels.

To evaluate the lateralization of alpha- and beta-band oscillatory activity during movement preparation, the right-hand movement condition was subtracted from the left-hand condition. To statistically compare both conditions, a paired t-test was computed on these differences, yielding a voxel-by-frequency matrix of t-statistics. To correct for multiple comparisons, a non-parametric permutation analysis was performed: the signs of the condition differences were randomly flipped, the paired t-test was recomputed, and the absolute maximum t-statistic was retained. This procedure was repeated 1000 times to construct a global null distribution. Empirical t-values exceeding the 95th percentile of this distribution (α = 0.05) were considered statistically significant.

## 3. Results

To assess the effectiveness of sBOSC in detecting oscillatory activity, we first applied the method to a series of simulations, analyzing the impact of different parameters on detection performance. Subsequently, sBOSC was applied to real resting-state MEG data to identify the brain’s natural frequencies at the single-voxel level, and these results were then compared with those obtained by Capilla et al. (2022). Finally, we tested sBOSC on a motor-task MEG dataset to assess whether it could detect the well-established modulations of alpha- and beta-band oscillations in the sensorimotor cortex (Pfurtscheller & Lopes da Silva, 1999; Salmelin & Hari, 1994).

### 3.1. Detection performance of sBOSC on simulated data

We applied sBOSC to a total of 405 simulations, each defined by a unique set of oscillatory activity and noise features. Oscillatory activity was simulated by a pure sine wave with different oscillatory frequencies (5, 10, and 20 Hz), number of cycles (3, 10, and 20 consecutive cycles), generator depth (65, 75, and 85 mm from the MEG sensors), and SNR (0.5, 1, and 1.5). The aperiodic component was simulated by adding the contribution of five generators of pink noise with semi-random locations to the signal (5 repetitions). After applying sBOSC, we computed hit and false positive rates for each set of parameters simulated.

On average, oscillatory episodes were accurately localized within a 1.5 cm radius of their simulated sources in 78% of all simulations. Among the remaining 22% of oscillatory episodes that were detected but mislocalized, 46% were localized within a 2.5 cm radius of their generators, while the other half decreased progressively with increasing distance. Only 15% of the mislocalized episodes (0.038% of all simulations) were found beyond 6.5 cm from their original location (Figure 3a). Moreover, more than half of the mislocalized episodes occurred in the lowest SNR scenario. In an ideal scenario (slowest frequency, longest duration, and highest SNR), the localization error is no longer observed, with all oscillatory episodes accurately localized within their sources.

**Figure 3.**
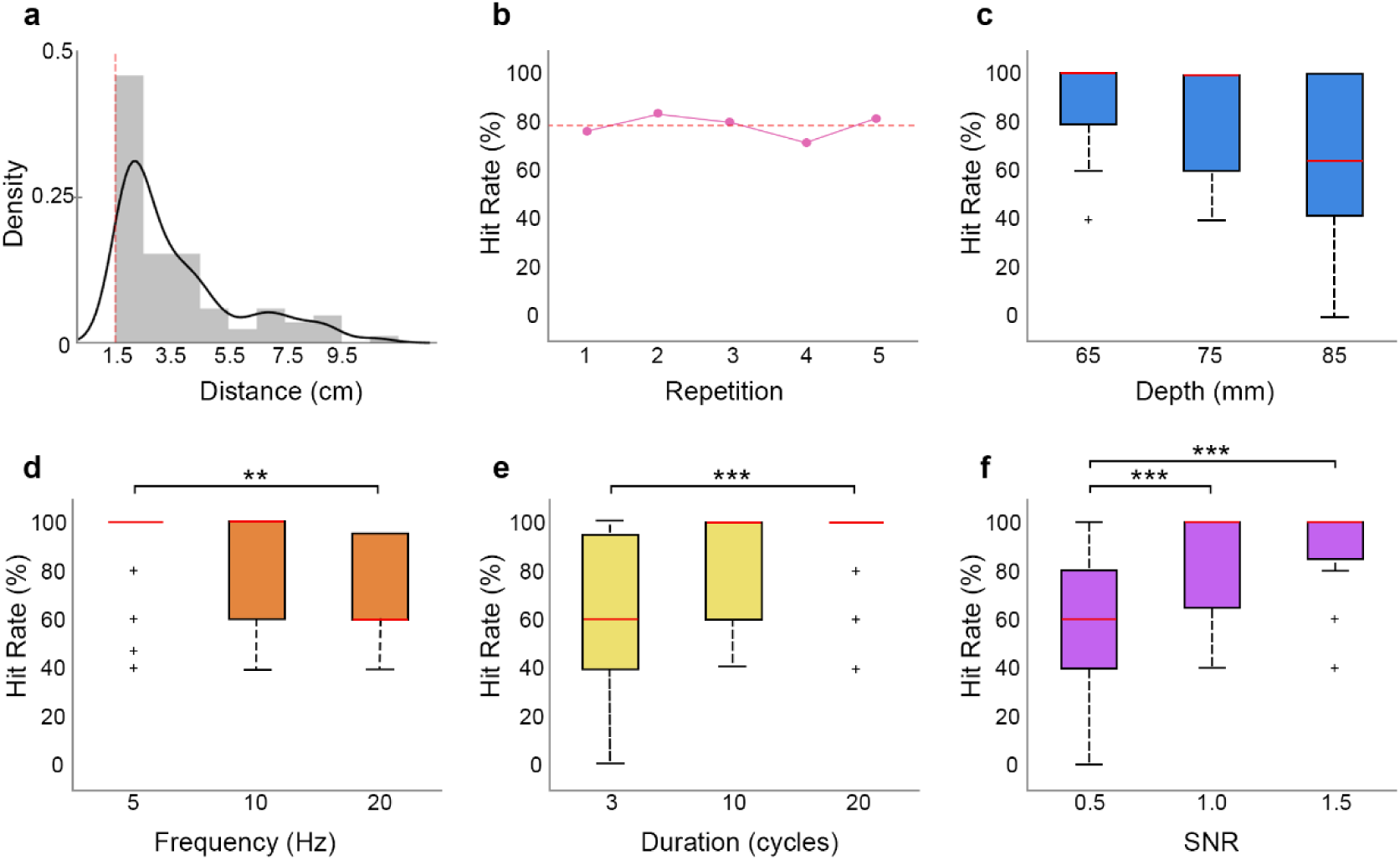
Influence of simulation parameters on sBOSC performance. **a.** Density distribution of detected episodes located more than 1.5 cm from the original source. Note that a substantial proportion of the errors occurred within 2.5 cm of the original source. **b.** Average hit rate in each repetition. Red dashed line indicates the mean across repetitions. **c-f.** Hit rate across the different parameter sets: depth from sensors, frequency, duration (number of cycles), and SNR. Red bars inside the boxes represent the median value. Significant post-hoc differences are indicated with asterisks (* p < .05, ** p <.01, *** p < .001).

The results of the Kruskal-Wallis test on the hit rate across repetitions showed that repetitions did not affect detection performance (H_(4,400)_ = 4.74, p = .315), thus ensuring that the semi-random location of the pink noise generators did not influence the results (Figure 3b). Additionally, distance from the signal source to the sensors (depth) was also not statistically significant (H_(2,78)_ = 5.37, p = .068) (Figure 3c). This was expected as SNR was adjusted independently of voxel depth. On the contrary, frequency (H_(2,78)_ = 11.36, p = .003; η_p_^2^ = .142), number of cycles (H_(2,78)_ = 15.54, p < .001; η_p_ ^2^ = .194), and SNR (H_(2,78)_ = 22.08, p < .001; η_p_ ^2^ = .276) significantly altered the hit rate detected with sBOSC. Specifically, lower frequencies (5 Hz), longer duration (20 cycles), and higher SNR (1.5) resulted in significantly greater hit rate compared to their counterparts (Figures 3d-f).

Since neither depth nor the location of the pink noise generators showed statistically significant effects, we averaged hit rate and false positive rate across these conditions to compute an overall performance of sBOSC. Figure 4 shows the resulting hit and false positive rates for each scenario. Correct detection of oscillatory episodes was on average 62% at the lowest SNR levels, increasing to 87% at neutral SNR and up to 90% at higher SNR levels. Importantly, false positive rate remained stable across different SNR levels, staying below 0.05% in all scenarios. Additional simulations conducted in the complete absence of true oscillatory signals yielded a comparable false positive rate (0.055% ± 0.0067 [mean ± sd), confirming the specificity of the method. Exceptionally low false positive rate values are due to the high number of negative cases (points without oscillations). Furthermore, we verified that using different window lengths for time-frequency decomposition (3-, 5-, or 7-cycle windows) did not significantly affect either hit rate (H_(2,24)_ = 2.81; p = .245) or false positive rate (H_(2,24)_ = .6; p = .742).

**Figure 4.**
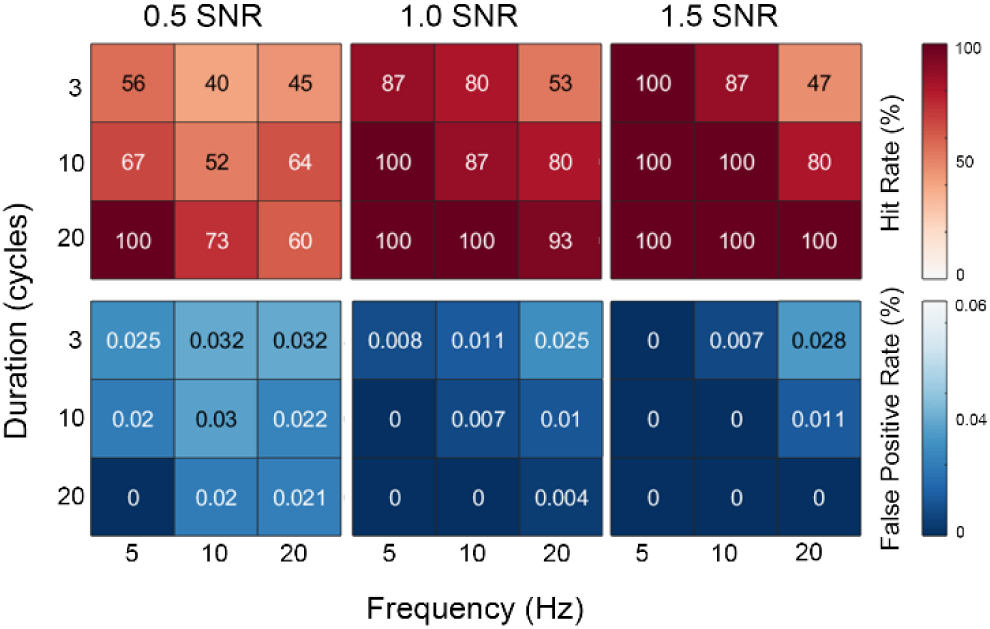
Detection performance of sBOSC in the simulated data. Hit rate (top panels) and false positive rate (bottom panels) are expressed as percentages across different scenarios, each defined by a combination of oscillatory frequency, number of cycles, and SNR levels. Parameter combinations resulting in the highest hit rate performance can be seen in warmer colors, while lower false positive rate are shown in cooler colors.

### 3.2. Characterization of the brain’s natural frequencies using sBOSC on resting-**state MEG data**

We applied sBOSC to a real set of resting-state MEG data. The supplementary material contains a video clip illustrating the detected oscillatory episodes for a representative participant (Supplementary Video 1). The same dataset had previously been used by Capilla et al. (2022) to identify the brain’s natural frequencies through k-means clustering of power spectra. Here, we aimed to test whether sBOSC could achieve similar results by detecting the most characteristic oscillatory frequency for each voxel. The comparison between the natural frequency map obtained using sBOSC and the original map is shown in Figure 5. Overall, the two maps exhibit a high degree of similarity, as indicated by a Pearson correlation coefficient of 0.634 (p<.001).

**Figure 5.**
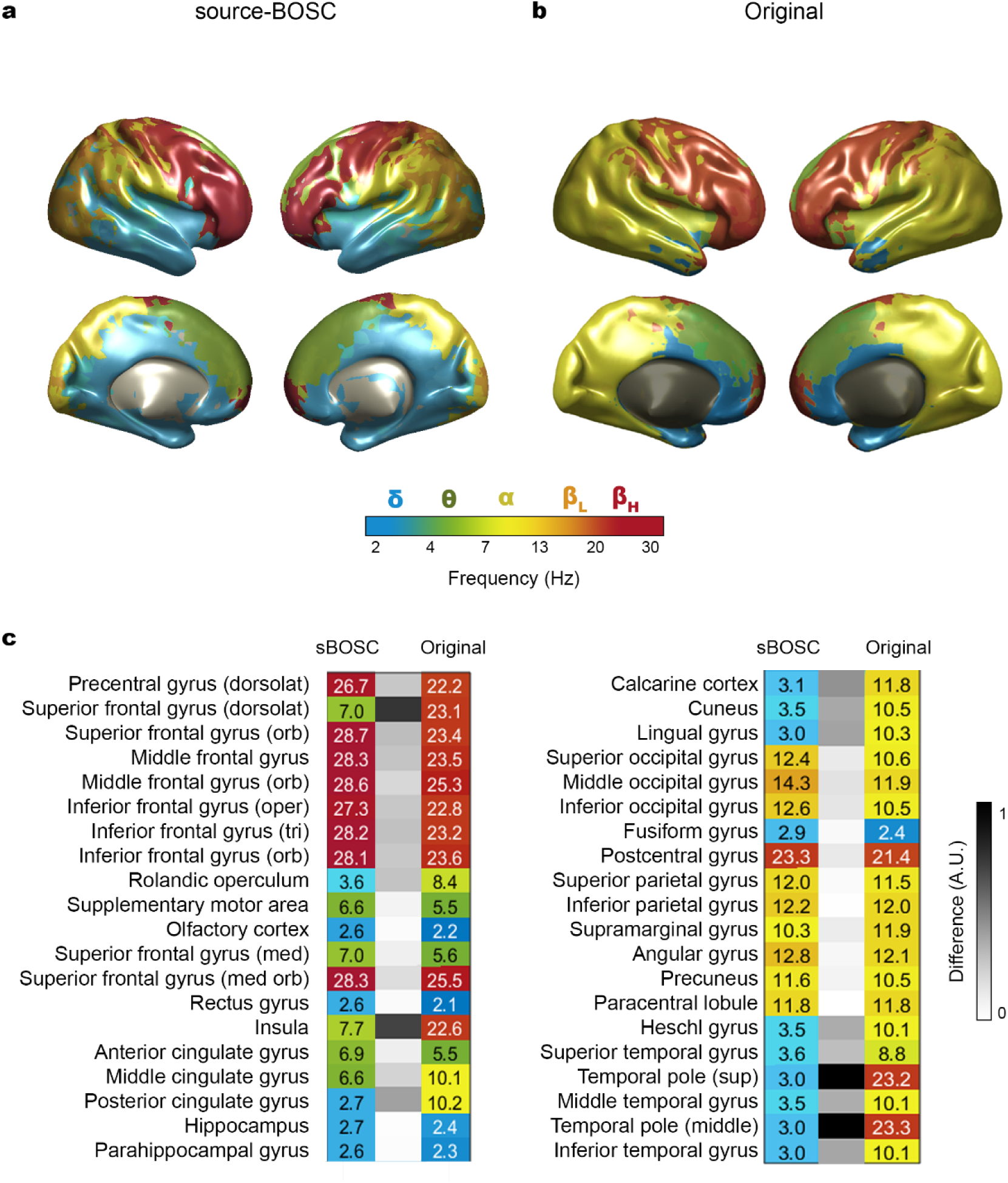
Brain map of natural frequencies derived using sBOSC. **a.** Natural frequencies extracted from the oscillatory episodes detected with sBOSC**. b.** Natural frequencies obtained Capilla et al. (2022) [Reproduced with permission]. **c.** Comparison of the natural frequencies across different AAL regions obtained with both methods. The color scale represents frequency (ranging from blue: delta band, to red: high-beta). White to black squares indicate the difference in the natural frequencies between the two methods for each region.

Both the voxel wise brain map and the distribution of natural frequencies across AAL regions exhibited a similar pattern between the two approaches. Delta frequency values (δ; 2-4 Hz) were characteristic of the orbitofrontal and temporal cortices, although they extended further into medial occipito-temporal regions with sBOSC, where the largest discrepancies were observed. Theta (θ; 4-8 Hz) represented the natural frequency of the medial frontal cortex. Alpha-band oscillations (α; 8-12 Hz) were distributed over parieto-occipital regions, although some areas of the parietal cortex showed a natural frequency within the low-beta instead of the alpha band using sBOSC. Finally, beta-band oscillations (β; 13-30 Hz) were characteristic of the sensorimotor and lateral prefrontal cortices.

### 3.3. Identification of lateralized alpha- and beta-band oscillatory episodes during a **motor preparation task**

Finally, to evaluate the method’s performance in a task context characterized by transient oscillations, we applied sBOSC to extract oscillatory episodes from epoched MEG data acquired during a motor task in the HCP dataset. Figure 6 displays the resulting t-statistics (corrected for multiple comparisons), contrasting left- versus right-hand motor preparation, for the alpha (8-12 Hz) and beta (18-30 Hz) bands. Positive t-values (red) indicate voxels with significantly lower oscillatory power (desynchronization) and shorter episode duration prior to right-hand movements compared to left-hand movements. Conversely, negative t-values (blue) indicate voxels with significantly lower power and shorter duration of oscillatory activity prior to left-hand movements compared to right-hand movements. As shown, both alpha- and beta-band activity exhibit stronger desynchronization over the contralateral sensorimotor cortex. Moreover, beta-band oscillatory sources are located slightly more anteriorly than alpha-band sources.

**Figure 6.**
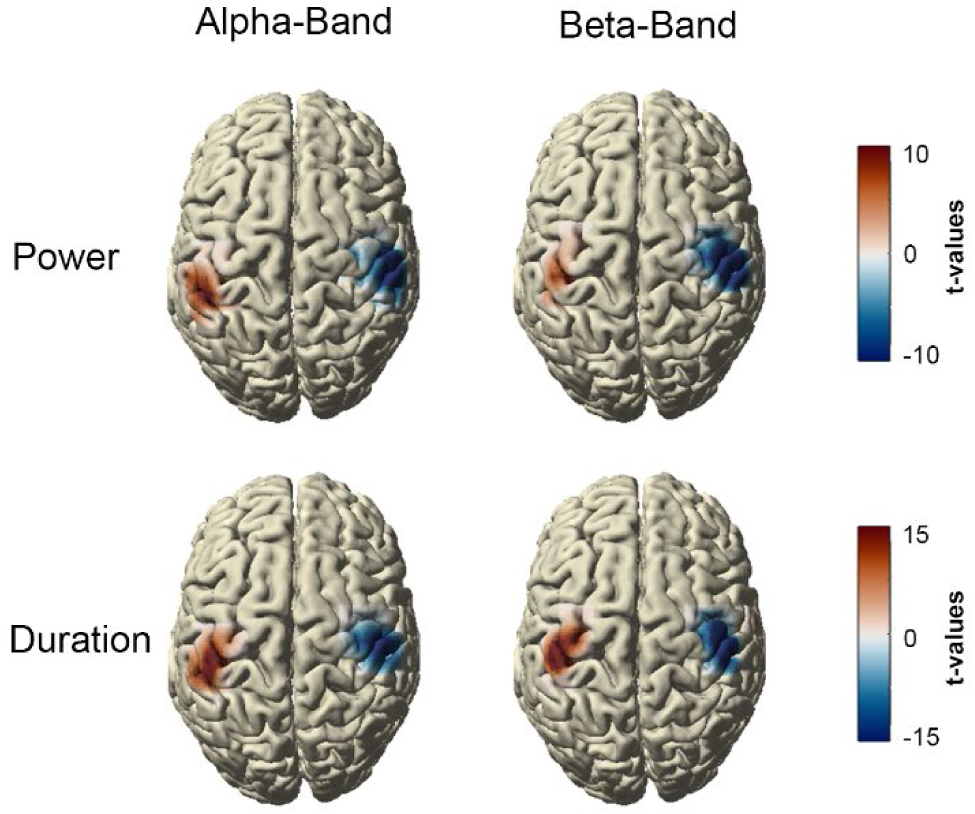
Lateralization of alpha- and beta-band oscillations during hand movement preparation. Brain maps show statistically significant differences (paired t-tests, corrected for multiple comparisons), contrasting left- versus right-hand movement preparation. Results are displayed for both the average power (top panels) and the duration (bottom panels) of oscillatory episodes detected with sBOSC within the alpha (8-12 Hz, left) and beta (18-30 Hz, right) bands. Positive t-values indicate regions where power and duration are lower prior to right-hand movements; negative t-values indicate regions where power and duration are lower prior to left-hand movements.

## 4. Discussion

In this study, we developed and validated a novel method for detecting oscillatory episodes at the source level in M/EEG data. This method, which we termed sBOSC (i.e., source-BOSC), builds upon the BOSC family of algorithms (Caplan et al., 2001; Kosciessa et al., 2020; Seymour et al., 2022; Whitten et al., 2011) and addresses two major limitations in current oscillation detection approaches. First, because true oscillations should manifest as spectral peaks rising above the aperiodic background (Donoghue et al., 2022; Gross, 2014; Lopes da Silva, 2013), our approach identifies local maxima in the frequency spectrum to distinguish broad-band high-power events from frequency-specific peaks. Secondly, most existing methods are restricted to single-channel signals in sensor space, whereas a comprehensive analysis of brain oscillations requires examining the neural sources generating that activity. With sBOSC, we have taken a step forward in the detection of oscillatory episodes by addressing both limitations.

Considered separately, the first methodological innovation—spectral peak detection—increases specificity by excluding points that exceed the amplitude threshold without forming true oscillatory peaks, thereby reducing false positives; this improvement applies equally at the sensor or source level and could potentially be integrated into existing algorithms. The second, and key, innovation—detecting oscillations directly in source space and focusing on voxels with local maxima—enhances spatial precision, avoids redundant detections, and improves interpretability. We first validated sBOSC through a series of simulations with different signal properties and then evaluated its performance on real resting-state and task-related MEG data. Importantly, although the validation was performed on MEG recordings, the algorithm is also applicable to EEG data.

### 4.1. sBOSC performance on simulated data

We evaluated the detection performance of sBOSC through a series of simulations defined by unique parameter combinations, included the location of the aperiodic component within the brain, the depth of the oscillatory generators, oscillation frequency, number of cycles, and SNR. We analyzed the algorithm’s performance across these different simulated scenarios.

The aperiodic component was semi-randomly generated at five locations within the brain, and thus this procedure was repeated five times for each unique parameter combination. No significant differences were observed between repetitions, thereby confirming that the location of the aperiodic generators did not affect sBOSC’s detection accuracy. Similarly, the location of the oscillatory sources did not influence detection performance either. Given the independent manipulation of the oscillation SNR regardless of the generator depth, we expected the location of the oscillator to have minimal or no effect.

We also evaluated the frequency and the number of cycles of the simulated oscillations. Significant differences emerged between the three frequencies analyzed, with the slowest frequency (5 Hz) being detected more often than the fastest frequency (20 Hz). Similarly, significant differences were found between the number of cycles, where the condition with the greatest number of cycles (20 cycles) yielded higher hit rate compared to the condition with the fewest (3 cycles). Both results can be explained by the total length of the oscillation, which increases with either a lower frequency or a higher number of complete cycles. Nonetheless, brain electrophysiological data contain oscillations from approximately 0.05 Hz up to 500 Hz (Buzsáki & Draguhn, 2004) that can last from short-lived transient bursts (Jones, 2016; Van Ede et al., 2018; Vidaurre et al., 2016) to several sustained cycles as in the case of posterior alpha oscillations. This implies that faster and/or shorter oscillatory activity may be challenging to detect with sBOSC. Moreover, sBOSC, along with other current oscillation detection methods, establishes a minimum requirement of three complete cycles for a given frequency to be considered an oscillatory episode. This threshold reduces the false categorization of random fluctuations in power as an oscillatory episode but comes at the cost of overlooking genuine transient oscillatory bursts that last less than three cycles. Thus, identifying oscillations occurring at shorter time scales remains a challenge. One possible approach for detecting transient brain states of brief duration is the use of Hidden Markov Models (HMM) (Quinn et al., 2019), although their direct relation to oscillatory neural activity is not yet fully understood.

Lastly, we assessed sBOSC performance across three different SNR scenarios, in which noise was higher (0.5), equal (1.0), or lower (1.5) relative to the signal amplitude. As expected, the algorithm yielded the highest detection accuracy (90%) when the signal amplitude was 50% greater than the noise. Even when signal and noise amplitudes were equal, accuracy remained high (87%) but declined (62%) in the most challenging condition, where noise exceeded the signal by 50%. It is worth noting that, in real MEG data, SNR is not constant but fluctuates across time, source location, and participants, resulting in mixed SNR conditions throughout the recording.

Overall, simulation results demonstrate that sBOSC is a robust algorithm for detecting source-level oscillatory episodes in multi-channel electrophysiological data, consistently achieving high detection rates and very low false positive rates. Detection performance is modulated by oscillation length and SNR, with longer-lasting oscillations and higher SNR yielding the most accurate results.

### 4.2. sBOSC performance in resting-state MEG data

After validating the method through simulations, sBOSC was subsequently applied to resting-state MEG data. The same participant dataset had previously been analyzed by Capilla et al. (2022) to extract a map of natural frequencies, which served as a benchmark for evaluating the results obtained with sBOSC. Once oscillatory episodes were identified, natural frequencies were computed as the frequency with the highest relative number of oscillatory episodes at each voxel. Group-level values were then calculated using the same bootstrapping procedure as Capilla et al. (2022) to ensure comparability.

Despite relying on different approaches to estimate natural frequencies (k-means clustering of power spectra versus oscillatory episode detection), both methods yielded consistent results, as evidenced by a strong voxel wise correlation (r = 0.634). Both approaches captured key oscillatory patterns. Delta frequencies were characteristic of the orbitofrontal and medial temporal regions, although sBOSC showed a broader extension over the temporal and occipital lobes compared to previous findings. This increased sensitivity to low-frequency oscillations may stem from explicitly removing the aperiodic component, which typically dominates the lower-frequency spectrum. In agreement with Capilla et al. (2022) (see also Arana et al., 2025), frequencies within the theta range consistently characterized the mid-frontal cortex. Alpha oscillations were the distinctive frequency of the posterior lobes, while parietal and lateral occipital regions were characterized by frequencies in the low-beta band. Finally, higher frequencies were also predominant in motor and prefrontal regions.

The frequency distribution identified with sBOSC is also in agreement with previously reported resting-state relative power maps derived from MEG (Keitel & Gross, 2016; Mellem et al., 2017; Niso et al., 2016) and iEEG data (Afnan et al., 2023; Kalamangalam et al., 2020). Characterizing resting-state brain activity presents specific challenges, particularly the strong dominance of the alpha rhythm due to its high amplitude and the effects of spatial leakage. As previously discussed, the data were spatially normalized to mitigate this dominance and enable direct comparison with the brain distribution of natural frequencies reported in Capilla et al. (2022). However, a given region may exhibit more than one characteristic oscillatory activity. Additional analyses, such as quantifying the number and duration of oscillatory episodes across canonical frequency bands, could also be performed using sBOSC, providing further insight into ongoing human neural oscillatory activity.

### 4.3. sBOSC performance in task-related MEG data

To further validate sBOSC on task-related data, we applied it to MEG recordings obtained during a motor task. Specifically, we examined the average power and duration of detected oscillatory episodes occurring prior to hand movement onset. Motor preparation is a well-characterized neurophysiological process; typically, sensorimotor regions contralateral to the movement exhibit a desynchronization of rhythmic activity in the alpha and beta ranges to prepare the motor system for action (Pfurtscheller & Lopes da Silva, 1999; Salmelin & Hari, 1994).

As expected, our analysis revealed significantly shorter oscillatory episodes with lower average power in these contralateral sensorimotor regions. Furthermore, while both alpha- and beta-band effects were localized to sensorimotor regions, beta-band activity was concentrated slightly more anteriorly. This finding is consistent with both classical studies (Pfurtscheller & Lopes da Silva, 1999; Salmelin & Hari, 1994) and more recent work (Stolk et al., 2019; Tzagarakis et al., 2015), which consistently report alpha-band oscillations localized to somatosensory cortex in the postcentral gyrus, whereas beta-band oscillations are predominantly found in the somatomotor cortex.

This final validation approach served a dual purpose. First, it provided a secondary validation step by successfully replicating gold-standard motor preparation effects using the oscillatory episodes identified with sBOSC. Second, it demonstrated the feasibility of applying sBOSC to single-trial, task-related data.

### 4.4. Comparison with other oscillation detection methods

A direct comparison of sBOSC with existing oscillation detection methods is limited due to the distinct scope of sBOSC. Methods such as the BOSC family (Caplan et al., 2001; Kosciessa et al., 2020; Seymour et al., 2022; Whitten et al., 2011) and PAPTO (Brady & Bardouille, 2022) are designed to identify oscillatory episodes in single-channel data. In contrast, sBOSC is specifically designed to detect the underlying sources of oscillations measured at the multi-channel level. Although existing methods could, in principle, be also applied to source-reconstructed multi-channel data, source leakage causes signals to spread across multiple voxels, thereby leading to an overestimation of oscillatory activity throughout the brain volume. sBOSC addresses this limitation by categorizing activity as an oscillatory episode only if the oscillation constitutes a spatial local maximum at that location. Ideally, this implies that locations where oscillatory episodes are identified correspond to genuine oscillators.

In practice, however, due to the combined influence of factors such as SNR, dipole orientation, and the inherent imprecision of source-reconstruction algorithms, the identified location cannot be guaranteed as the true oscillatory generator (Brunner et al., 2016; Jonmohamadi et al., 2014; Van Den Broek et al., 1998; Van Veen et al., 1997; Westner et al., 2023). In fact, as demonstrated in our simulations, oscillatory episodes identified more than 1.5 cm away from the actual generator were treated as misses, even though their frequency and timing were correctly recovered. This occurred in 22% of the total simulations. While sBOSC is designed to detect oscillatory episodes originating from their true underlying sources, caution is warranted when inferring that any particular episode location corresponds to a genuine oscillator.

### 4.5. Applicability and remaining challenges

Several advantages arise from detecting oscillatory episodes directly, rather than interpreting relative power changes, as it is often done in M/EEG research by subtracting a baseline window presumed to containing no relevant activity (Gross, 2014; Gyurkovics et al., 2022).

First, sBOSC allows for the effective analysis of electrophysiological signals in situations where baseline windows are not available, such as resting-state recordings or experimental tasks with short intertrial intervals, while still accounting for the pervasive aperiodic activity. To extract the aperiodic component in simulations as well as in the resting state data, 20-second segments were used as a trade-off between computational efficiency and sufficient sensitivity to track dynamic fluctuations in aperiodic activity over time (Gao et al., 2017). However, this segment length can be adapted by the user, for example, when handling epoched data.

Second, analyzing relative power changes in the power spectrum may lead to misleading interpretations of oscillatory brain activity, as these changes can reflect variations in the aperiodic component rather than genuine oscillatory dynamics (Donoghue et al., 2022). By focusing only on activity that exhibits a local maximum above the aperiodic background, sBOSC ensures that the analysis targets true oscillatory phenomena, thereby avoiding misinterpretations driven by non-oscillatory power fluctuations.

Third, detecting oscillatory episodes directly in source space might substantially simplify connectivity analyses, which often explore relationships across various signal features (e.g., amplitude, phase, or phase-amplitude coupling) across multiple frequencies and brain regions. Focusing on segments where oscillatory episodes are identified at specific locations would help narrow down the vast number of potential hypotheses, thereby reducing the computational load and increasing the interpretability of connectivity results.

While sBOSC offers potential benefits, it also has some limitations. First, it is tuned to detect episodes with a minimum duration of three cycles. This conservative threshold serves to filter out noisy transients but may also exclude genuine short-lived oscillatory bursts (Jones, 2016; Van Ede et al., 2018; Vidaurre et al., 2016). However, this parameter can be adjusted by the user to identify shorter episodes if needed at the cost of increasing the number of false positives.

A second problem arises when interpreting the location of identified episodes as their true generators. Although most of our simulations successfully recovered the original sources of the oscillations, this limitation is inherent to the ill-posed nature of source-reconstruction algorithms (Brunner et al., 2016; Jonmohamadi et al., 2014; Van Den Broek et al., 1998; Van Veen et al., 1997). Accurate identification of oscillatory episodes depends not only on sBOSC’s detection performance but also con the precision of beamformer reconstruction. Consequently, an oscillatory episode that is correctly detected by sBOSC may still be considered a miss if the beamformer’s reconstructed location deviates from the true source. This limitation is likely to be more pronounced when applying sBOSC to EEG rather than MEG signals, as volume conduction effects reduce the precision of source reconstruction.

## 5. Conclusion

In this work, we introduce sBOSC, an algorithm specifically designed to detect oscillatory episodes in M/EEG data at the source level. We validated the method using both simulated and real datasets. This algorithm extends the BOSC family of methods by supporting multi-channel data and ensuring the presence of oscillatory activity above the aperiodic background. sBOSC offers a complementary approach for studying oscillatory dynamics at the source level, enabling the identification of non-averaged oscillatory events over time and opening new avenues for measuring brain connectivity.

## Code and availability

The full code required to reproduce the method, analyses, and figures in this work is openly available at https://github.com/necog-UAM/.

## Author Contributions

**Enrique Stern**: Conceptualization, Data curation, Formal analysis, Methodology, Software, Visualization, Writing– original draft preparation; **Guiomar Niso**: Conceptualization, Writing – Review & editing; **Almudena Capilla**: Conceptualization, Data curation, Funding acquisition, Methodology, Writing – original draft preparation.

## Acknowledgements

We are thankful to the contributors of Open MEG Archive (OMEGA) for providing the publicly available dataset used in this work.

This work was supported by Ministerio de Ciencia e Innovación / Agencia Estatal de Investigación, Spain / FEDER/FSE+, UE (MCIN/AEI/10.13039/501100011033/FEDER/FSE+, UE; grants PRE2022-101613 to ES, RYC2021-033763-I and PID2023-150034OA-I00 to GN, and PID2021-125841NB-I00 and PID2024-161032NB-I00 to AC).

